# Rapid evolution of *Wolbachia* genomes in mosquito cell culture

**DOI:** 10.1101/2023.09.20.558649

**Authors:** Julien Martinez, Steve P. Sinkins

## Abstract

*Wolbachia* bacterial symbionts are widespread across arthropods where they cause reproductive manipulations and/or confer fitness benefits such as protection against viral pathogens. Their self-spreading ability coupled with their antiviral effect has been harnessed in health programmes to curb the transmission of dengue virus. Comparative genomics of *Wolbachia* strains has been a useful tool to understand the general trends in the evolution of the symbiont genome; however, short-term evolutionary processes occurring within hosts remain poorly explored. Understanding these short-term dynamics is necessary to provide a more complete picture of *Wolbachia* evolution and will inform ongoing *Wolbachia*-based disease control interventions. Here we generated six new mosquito cell lines by introducing a range of *Wolbachia* strains from *Drosophila* into the symbiont-free *Aedes albopictus* Aa23 cell line. Following transinfection, we tracked temporal variation in *Wolbachia* density and identified *de novo* mutations through re-sequencing of the symbiont genome. Several mutations were associated with major shifts in bacterial density. Moreover, signs of parallel evolution across cell lines, combined with an excess of non-synonymous mutations, indicate that *Wolbachia* evolution in cell culture is dominated by selective processes rather than genetic drift. Our results also provide new candidate genes likely to be involved in symbiont density regulation. Altogether, our study demonstrates that cell culture is a valuable tool to investigate symbiont short-term evolution, identify the genetic basis of bacterial density variation and for the generation of new higher-density variants for use in control programmes.

## Introduction

*Wolbachia* is an intracellular and maternally-inherited bacterial symbiont found in a wide range of arthropods and some filarial nematodes (Zug and Hammerstein 2012). Its maternal transmission is ensured by the symbiont’s ability to colonize the female germline and be passed on to the eggs. *Wolbachia* has evolved various kind of reproductive manipulations, such as cytoplasmic incompatibility, that promote the production of *Wolbachia*-carrying hosts in the population (Werren et al. 2008). In addition to the germline, *Wolbachia* is often found in somatic tissues which can induce large effects on the host fitness (Pietri et al. 2016). In particular, many strains of the symbiont inhibit the replication of viruses, a property that has been harnessed by health programmes around the world to control the spread of dengue virus (Teixeira et al. 2008; Moreira et al. 2009; Bian et al. 2010; Hoffmann et al. 2011; Walker et al. 2011; Martinez et al. 2014; Ant et al. 2018; Nazni et al. 2019; Utarini et al. 2021). Higher *Wolbachia* densities within host cells generally provide stronger virus inhibition but can also induce substantial fitness costs by decreasing the host lifespan or fecundity (Chrostek et al. 2013; Martinez et al. 2014; Duarte et al. 2021).

While symbiont density regulation is central to the evolution of host-*Wolbachia* associations and the efficacy of *Wolbachia*-based disease control, the underlying genetic basis and mechanisms remain poorly understood. *Wolbachia* lives in the host cell cytoplasm within host-derived membranes and this intracellular lifestyle has hampered the development of genetic tools to manipulate and study the symbiont’s genome (Porter and Sullivan 2023). Therefore, *Wolbachia* research has been largely reliant on the characterization of natural genetic variation and comparative genomics to identify density-associated candidate genes (Chrostek et al. 2013; Baião et al. 2021; Hague et al. 2021; Martinez et al. 2022). This approach was successful in identifying a 21 kb density-regulating genomic region called Octomom in the pathogenic strain *w*MelPop. *w*MelPop overproliferates at high temperatures in *Drosophila melanogaster*, causing severe reductions in host longevity (Min and Benzer 1997). This overproliferating phenotype was found to be associated with a higher Octomom copy number in *w*MelPop compared to the closely-related and lower-density strain *w*MelCS which only harbours one copy of the region (Chrostek et al. 2013; Woolfit et al. 2013). In *w*MelPop, Octomom copy numbers also vary between host generations, as well as with the age of the fly host, with more copies being associated with a higher *w*MelPop density, pathogenicity and levels of virus inhibition (Chrostek and Teixeira 2015; Chrostek and Teixeira 2018; Bénard et al. 2021; Monnin et al. 2021). Such genomic instability is facilitated by the presence of repeat elements flanking the Octomom region. Surprisingly, the complete loss of Octomom has similar effects to the amplification of the region by also increasing symbiont density, pathogenicity and virus inhibition compared to a single-copy variant (Duarte et al. 2021), suggesting complex interactions between the Octomom copy number and *Wolbachia* proliferation.

Symbiont density regulation is most likely determined by several biological pathways involving many other genes yet to be characterized. Cases like the Octomom region in which *Wolbachia* strains maintained in the lab only vary at a limited number of loci are rare. Thus, comparative genomics approaches tend to generate long lists of candidate genes which association with density often remain hypothetical (Baião et al. 2021; Martinez et al. 2022). A complementary approach is to track genomic changes occurring over short timescales. However, re-sequencing of *Wolbachia* genomes following the artificial transfer of the symbiont between host species or its introduction in the field for disease control revealed a relative stability of *Wolbachia* genomes, even over the course of several years to more than a decade (Huang et al. 2020; Morrow et al. 2020; Baião et al. 2021; Martinez et al. 2022). In contrast to insect studies, there is scarce evidence that *Wolbachia* genomes may evolve rapidly when the symbiont is maintained in insect cell culture. Woolfit et al. (2013) observed a rapid burst of genomic changes during the passaging of *w*MelPop in mosquito cell lines but detected no further mutations after transinfection into mosquitoes. These mutations included the complete loss of the Octomom region, the insertion of a transposable element and three other non-synonymous mutations (Woolfit et al. 2013). Finally, in the closely-related *w*Mel strain, major genomic changes were also detected in a cell line derived from *D. melanogaster*, with a complete loss of the Octomom region as well as a ∼3-4 fold amplification of a 35 kb region (Grobler et al. 2018). These studies suggest that *Wolbachia* may evolve more rapidly in cell culture, perhaps due to different population dynamics and selective pressures in this environment.

Here we investigated the evolution of several *Wolbachia* strains from *Drosophila* after transinfection and passaging into *Aedes albopictus* mosquito cell lines. We then examined the dynamics of occurrence and spread of the identified mutations and explored their potential association with changes in symbiont density. Our results support the hypothesis that *Wolbachia* genomes evolve more rapidly in a cell culture environment with important consequences on symbiont density and implications for *Wolbachia*-based control programmes.

## Results

### New *Wolbachia* strains stably colonize mosquito cell lines

In total six symbiont strains belonging to the arthropod-specific *Wolbachia* supergroup A were used to generate new *Wolbachia*-infected mosquito cell lines. The symbiont strains originated from various *Drosophila* species and were previously maintained in the same inbred line of *D. simulans* (STCP background) for six years or longer depending on the strain (Table 1). The *D. simulans* lines were used as a source to infect *Wolbachia*-free *Ae. albopictus* Aa23 cells from fly embryos (see methods). While all six *Wolbachia* strains successfully colonized mosquito cells (Figure 1A), bacterial densities were highly variable across time, with short-term fluctuations and, in some cases, sustainable shifts in bacterial density, as observed in *w*Tri-2-infected cells (Figure 1B).

**Figure 1.**
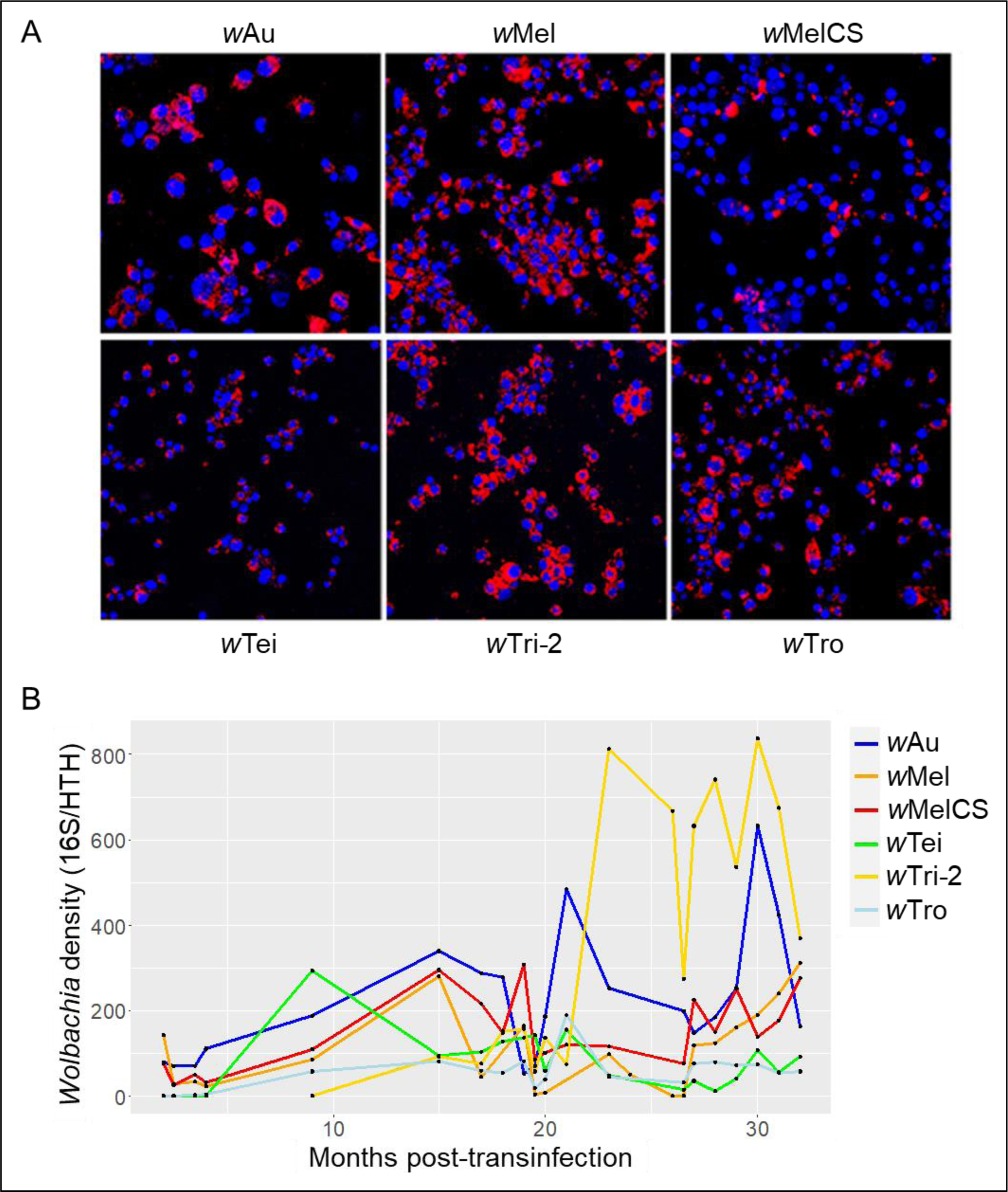
Long-term dynamics of *Wolbachia* density in newly-created Aa23 cell lines. (A) Fluorescent *in situ* hybridization imaging of Aa23 cell lines conducted >2 months post-transinfection, showing *Wolbachia* (red, 16S rRNA probe) and host nuclei (blue, DAPI). (B) *Wolbachia* density per cell line across passages following transinfection. The *w*Tri-2-infected cell line was generated five months after the other cell lines.

**Table 1.** Origin of *Wolbachia* strains used for transinfection of Aa23 mosquito cells.

| *Wolbachia* strain | Reference genome accession | Native host | Time spent in the *D. simulans* STCP line | References | Mean depth after re-sequencing |
| --- | --- | --- | --- | --- | --- |
| wAu | CP069055.1 | *D. simulans* | > 12 years | (Zabalou et al. 2008) | 677x<sup>a</sup> |
| wMel | CP046925.1 | *D. melanogaster* | > 16 years | (Poinsot et al. 1998) | 278x<sup>a</sup>, 135x<sup>b</sup> |
| wMelCS | CP046924.1 | *D. melanogaster* | 6 years | (Martinez et al. 2014) | 348x<sup>a</sup> |
| wTei | CP069052.1 | *D. teissieri* | > 12 years | (Zabalou et al. 2008) | 299x<sup>a</sup> |
| wTri-2 | GCA_014129515.1 | *D. triauraria* | 6 years | (Martinez et al. 2014) | 494x<sup>c</sup>, 206x<sup>d</sup> |
| wTro | GCA_014129525.1 | *D. tropicalis* | 6 years | (Martinez et al. 2014) | 77x<sup>a</sup> |

<sup>a, b, c, d</sup> re-sequencing after 17 (<sup>a</sup>), 27(<sup>b</sup>), 12(<sup>c</sup>) and 22(<sup>d</sup>) months post-transinfection.

Part of these fluctuations in *Wolbachia* density may be attributable to complex population dynamics or variation in other factors in the cell culture environment. Remarkably, when replicate cell lines were derived from the same ancestral population of cells, fluctuations in symbiont density were highly correlated between replicates, despite the fact that they were maintained as separate lineages for 150 days (Figure S1). However, density fluctuations did not appear to be strongly correlated between *Wolbachia* strains, suggesting a strain-dependent phenomenon (Figure S1). *Wolbachia* densities tended to be influenced by the host cell number seeded upon each passage to new flasks, with more cells leading to higher symbiont densities (Figure S2). High *Wolbachia* densities also tended to be associated with a reduced growth rate of host cells, indicating that high-density *Wolbachia* strains could be selectively disadvantaged under the hypothesis that *Wolbachia* is purely vertically-transmitted upon host cell division (Figure S2). However, the symbiont fitness could be partially decoupled from the host fitness if the symbiont can move horizontally between cells. We tested this hypothesis by co-culturing cells carrying one of the Supergroup A strains with cells harbouring the Supergroup B *w*AlbB strain native of *Ae. albopictus*. After 3 months, co-cultured *Wolbachia* strains were found within the same host cells, showing that, in addition to vertical inheritance, horizontal transmission of the symbiont is at play in this cell culture model (Figure S3).

### *Wolbachia* genomes accumulate non-synonymous mutations in cell culture

Besides the effects of population dynamics or environmental fluctuations, some changes in *Wolbachia* density could have been caused by evolutionary changes in the genome of the symbiont and/or the host following transinfection. In order to identify *de novo* mutations in the symbiont, we re-sequenced the genomes of the transinfected *Wolbachia* strains (Figure 2). Paired-end Illumina reads were mapped onto their respective reference genomes generated previously from their native *Drosophila* species, with mean sequencing depths ranging from 77x to 677x per genome (Table 1, Figure S4). Sequencing depth variation along the genomes were visually inspected and a frequency threshold of 0.9 was used to detect Single Nucleotide Polymorphisms (SNPs) and small indels. We found no difference between the *w*Au genome from our cell line and the reference sequence from its original *Drosophila* host. Three SNPs that were detected both using reads from our *w*Au-infected Aa23 cell line and those used to assemble the *w*Au reference genome (SRA accession: SRR13529481) were all located in repeated intergenic regions and showed sign of within-sample polymorphism indicating there are most likely sequencing errors. On the contrary, the five other strains all differed from their reference sequence by the presence of SNPs, small indels, large deletions or gene amplifications (Figure 2). We found no additional variation using a threshold of 0.2 to detect genetic variants segregating at low to intermediate frequencies.

**Figure 2.**
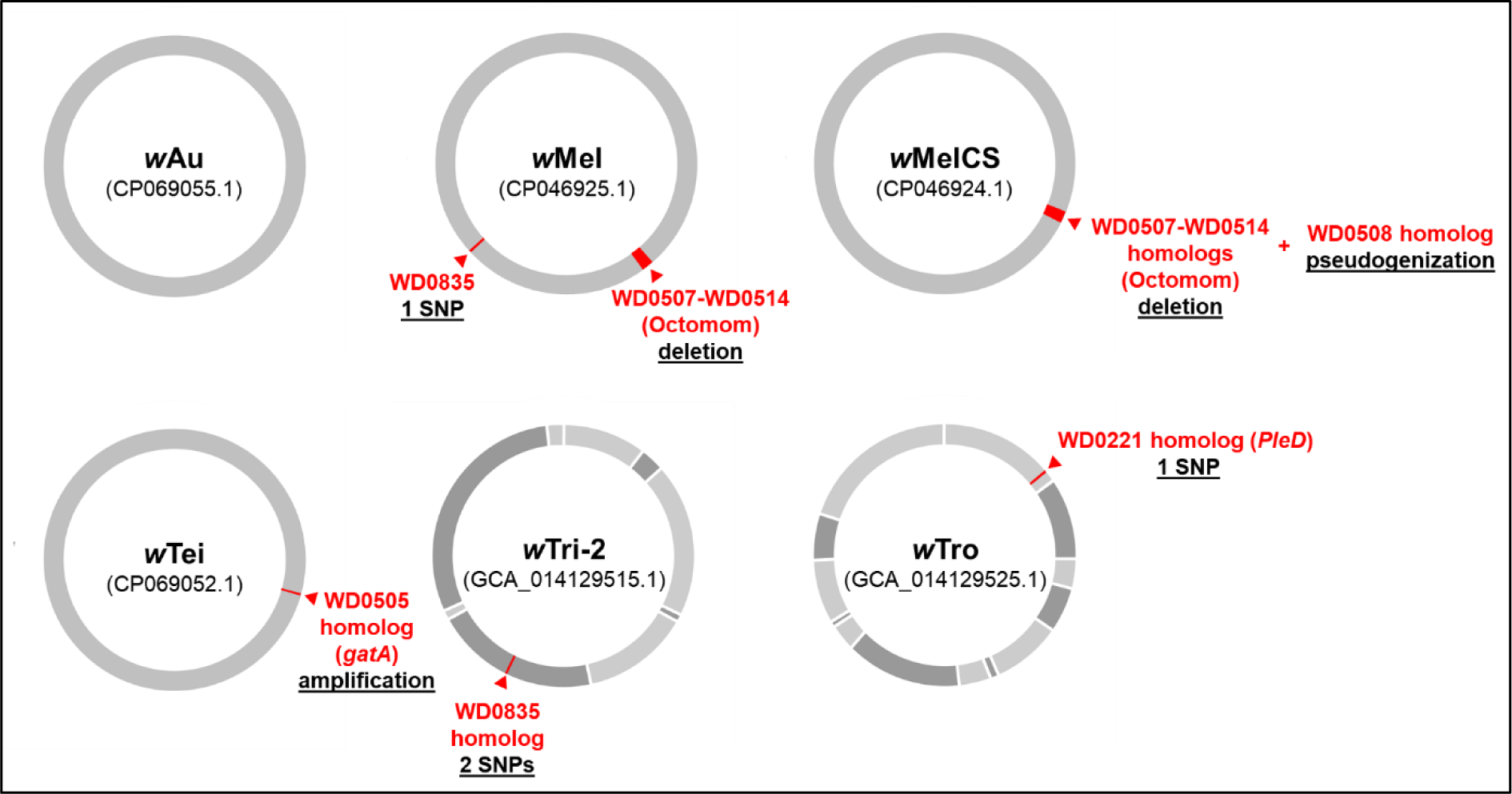
Summary of *de novo* mutations detected in each *Wolbachia* genomes (Genbank assembly accessions used as a reference in parentheses).

Multiple SNPs and short indels were found in the case *w*Tro and *w*Tri-2 (Table S1). However, the reference genomes for these two strains are incomplete and fragmented, which could produce false variants, in particular around misassembled repeat regions. Many variant sites in *w*Tro and *w*Tri-2 were indeed located within or in the flanking regions of repeat elements (transposase, reverse-transcriptase) and visual inspection of the mapped reads often revealed clear signs of polymorphism at those positions within samples (Table S1). In addition, a majority of these SNPs were also detected with the original sequencing data used to assemble the *w*Tro and *w*Tri-2 genomes (SRA accessions: SRR10988561, SRR10988562), suggesting that these variant sites are most likely false-positives due to misassembled regions (Table S1). Excluding these dubious variants, 2 sites were retained as potentially genuine variants in *w*Tro, one of which is located in a repeated intergenic region and could not be validated by Sanger sequencing. The other variant in *w*Tro is a SNP inducing a single amino acid change in a gene encoding a homolog of the *w*Mel gene WD0221 and was confirmed as genuine by Sanger sequencing (Figure S5). In the case of *w*Tri-2, after filtering out dubious variants, we detected 3 variants, of which one is located in a repeated intergenic region and could not be confirmed, and two are non-synonymous SNPs in a gene encoding a hypothetical protein homolog of the wMel gene WD0835.

Apart from *w*Au, *w*Tri-2 and *w*Tro, all other strains only differed from their reference genomes by one to two mutations. The absence of these mutations in the *D. simulans* lines used as donors was also confirmed by Sanger sequencing or quantitative PCR (qPCR) in the case of copy number variation, demonstrating that these mutations were acquired after transfer to the Aa23 cell line (Figure S5). All mutations but one induced non-synonymous changes and are described in detail thereafter. The only SNP located in an intergenic region in *w*Tei was also detected using the raw reads generated to assemble its reference genome (SRA accession: SRR13529477) and showed sign of within-sample polymorphism (Table S1). The intergenic region overlapping the SNP is highly repeated in the *w*Tei genome and our attempts to confirm it with Sanger sequencing from the donor flies and from Aa23 cells were both unsuccessful. Therefore, we considered this intergenic SNP as a sequencing error in the reference genome.

### Repeated loss of the density-regulating Octomom region

At the first round of sequencing (17 months post-transinfection), a ∼50% drop in sequencing depth overlapping the Octomom region (genes WD0507-WD0514) was observed in *w*MelCS, indicating the partial loss of this genomic region (Figure 3A, Figure S4). This was not observed at the same timepoint in the closely-related strain *w*Mel, which also harbours the region, but in which the sequencing depth was evenly distributed along the genome (Figure S4). We confirmed the presence of Octomom-deleted variants in the *w*MelCS-infected cell line by PCR, using primers designed in the flanking regions of Octomom (Figure 3B). In the donor flies, the Octomom region was present but the deleted variant was not detectable (Figure 3B). In Aa23 cells, Octomom-deleted variants of *w*MelCS were detected as early as 3.5 months post-transinfection and persisted until the end of the experiment. However, the presence of the Octomom gene WD0513 in all *w*MelCS-infected samples indicated the region was never completely eliminated in this cell line (Figure 3B). A similar phenomenon was observed in the *w*Mel-infected Aa23 cell line, with Octomom-deleted variants being first detected by endpoint PCR 9 months post-transinfection (Figure 3B). In the case of wMel, however, we did not detect the presence of Octomom after month 23, indicating the complete loss of the region. This was confirmed by re-sequencing the *w*Mel genome 27 months post-transinfection, with no more reads mapping to the region (Figure 3A and Figure S4).

**Figure 3.**
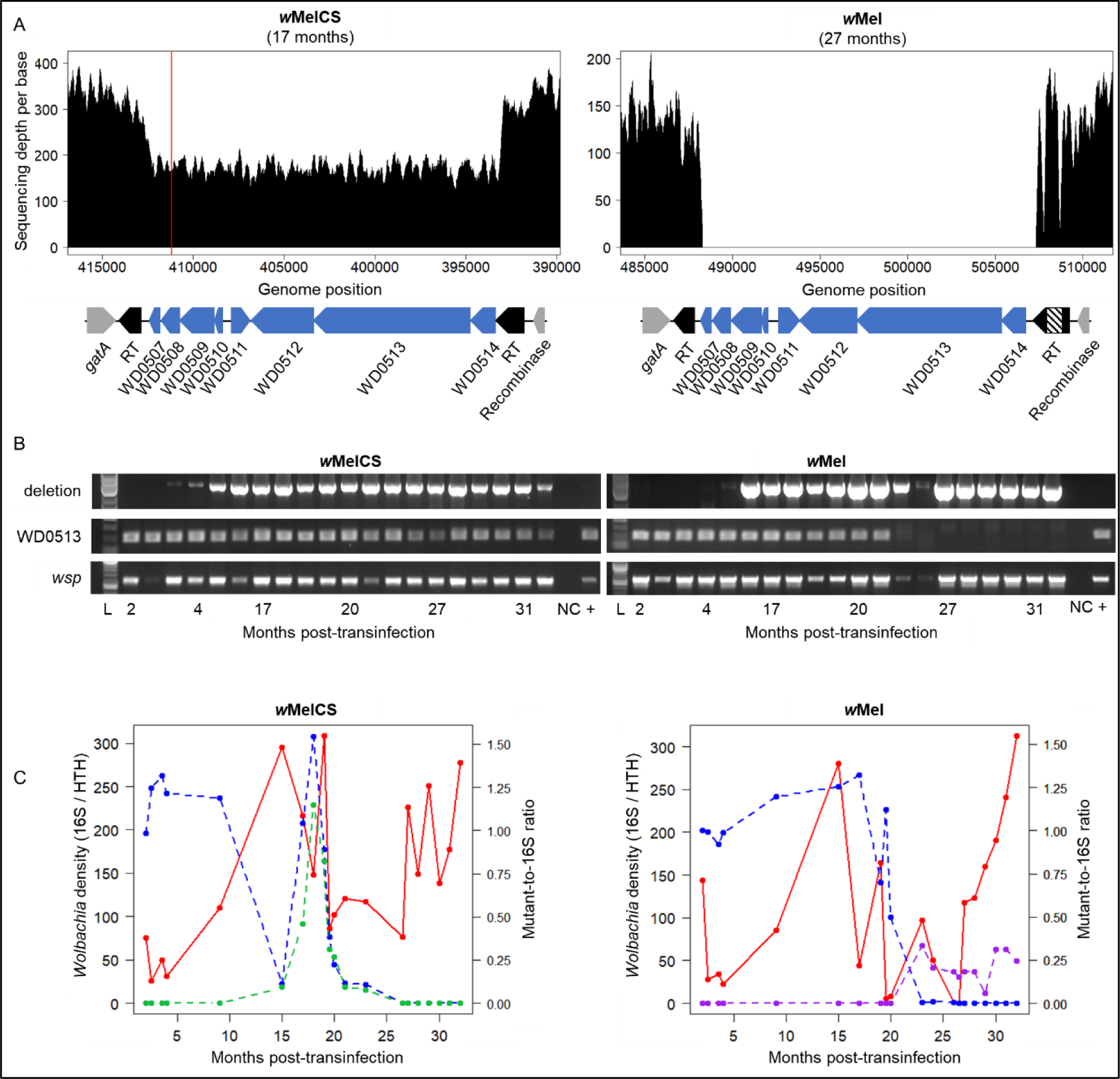
Evolution of the Octomom region. (A) Variation in sequencing depth on Octomom (blue boxes) and flanking regions in *w*MelCS- and *w*Mel-infected Aa23 cell lines. The vertical red line indicates the position of a SNP in the WD0508 homolog of *w*MelCS. RT: reverse-transcriptase. The dashed box in *w*Mel Octomom region indicates and transposase insertion disrupting the reverse-transcriptase. (B) Gel electrophoresis of PCR products for detecting Octomom variants in samples collected at different timepoints following transinfection. The *wsp* gene was used as a positive control for *Wolbachia* presence. The *w*MelCS- and *w*Mel-infected *D. simulans* STCP line were used as positive controls (+). NC: H_2_O negative control. L: DNA ladder. (C) Variation in *Wolbachia* density (red) and Octomom mean copy number (blue) per sample across time as measured by qPCR. Green: estimated frequency of the WD0508 SNP in *w*MelCS. Purple: estimated frequency of the WD0835 SNP in *w*Mel.

In order to obtain a better resolution of the changes in Octomom, we then monitored the relative copy number of the region throughout the experiment using qPCR. Similar to the Octomom copy number in the original *Drosophila* hosts, Octomom in *w*Mel- and *w*MelCS-infected Aa23 cells started at around one copy per genome following transinfection (Figure 3C). In *w*MelCS, copy numbers decreased to 0.11 Octomom copies per genome on average at month 15, increased again to around 1.5 copies and finally decreased to around 0 copy after 25 months. Together with the endpoint PCR data, this indicates that Octomom-carrying *w*MelCS genomes persisted at extremely low frequency towards the end of the experiment. When Octomom copy numbers were below 1, *w*MelCS densities tended to be higher than when copy numbers fluctuated around 1, similar to what was previously observed when the region is absent in *Drosophila* hosts (Duarte et al. 2021). Interestingly, the rebound and subsequent decrease in Octomom copy number in *w*MelCS between month 15 and 21 was tightly associated with the spread of an ORF-disrupting mutation in gene WD0508, a putative transcription regulator found in the Octomom region (Figure 3A and 3C, Table S1).

In *w*Mel, Octomom copy numbers only started to decrease from month 17 onwards, until the region was completely lost (Figure 3B and 3C). However, unlike *w*MelCS, *w*Mel densities did not appear to follow the variation in Octomom copy number. Moreover, a non-synonymous mutation that was detected through genome sequencing at month 27 in the WD0835 gene (Figure 2 and 3C; Table S1) appeared concomitantly with the progressive loss of the Octomom region and was then maintained at intermediate frequencies in the cell line. The evolution of WD0835 and its homologs is discussed in more details below.

In order to test whether the Octomom deletion spreads through the cell culture randomly under the sole effect of genetic drift or whether selection is at play, we generated six additional *Wolbachia*-infected cell lines from flies for *w*MelCS, as well as for the closely-related strain *w*MelPop in which Octomom is unstable, with copy number varying above one in *Drosophila* hosts. For both strains, the replicate cell lines were maintained separately and passaged in parallel. To minimize stochastic effects that could affect variation in both Octomom copies and *Wolbachia* density, the cell lines were split at regular intervals (every ten days) for about 400 days. In these new cell lines, Octomom started to be lost after about 200 days in most replicates (Figure 4A). Concomitantly with the loss of Octomom, we observed a sustainable increase in *Wolbachia* density, suggesting that, as seen in flies, Octomom deletion positively affects symbiont density (Figure 4A). Although rebounds in the Octomom copy number was observed for two *w*MelCS replicates, the overall decrease in Octomom copy number indicates that Octomom-deleted *Wolbachia* genomes may be at a selective advantage when maintained in cell culture. While Octomom copy numbers converged towards 0 copies in most replicates, endpoint PCR showed that the region generally persisted at low frequency except in two replicates in which the region was undetectable after 410 days of passaging (Figure 4B). Interestingly, the drop in Octomom copy number was preceded by an increase above 1 copy in most replicates. This could be due to a mutational bias where the deletion of the region is more likely to occur when the Octomom copy number is higher than one. Finally, the relative synchronicity in the loss of Octomom across replicates suggests that some unknown environmental factor fluctuating over time may be driving this selective advantage.

**Figure 4.**
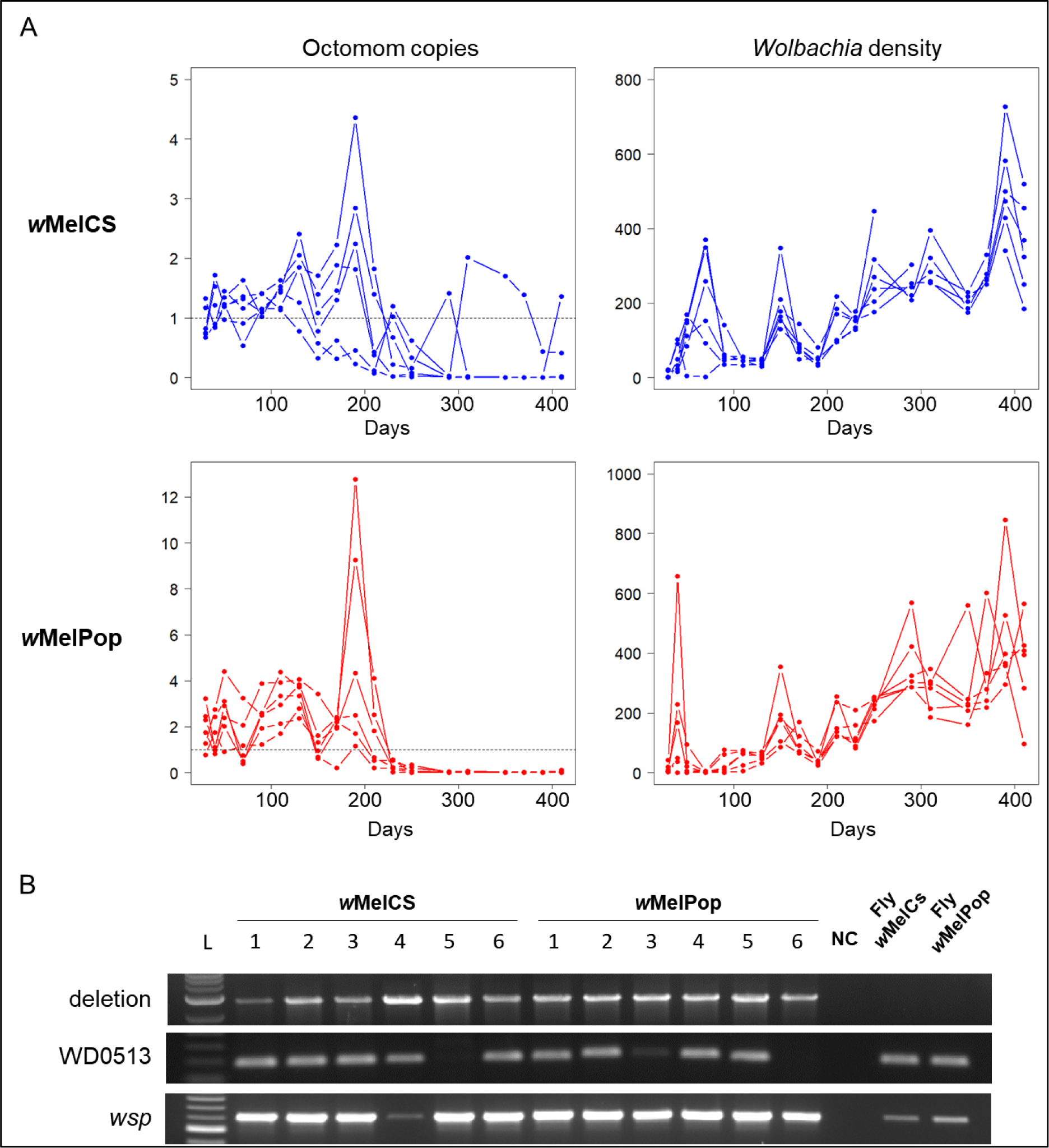
Repeated loss of the Octomom region in replicate cell lines. (A) Relative Octomom copy number across time estimated by the qPCR amplification of the WD0513 homolog (left panels) in *w*MelCS- (top) and *w*MelPop-infected (bottom) cell lines. Right panels show the corresponding *Wolbachia* densities. (B) Gel electrophoresis of PCR products for detecting Octomom variants in samples collected 410 days post-transinfection (290 days for *w*MelCS replicate 4 which was lost after that). The *wsp* gene was used as a positive control for *Wolbachia* presence. L: DNA ladder. NC: H_2_O negative control.

### Parallel evolution of WD0835 homologs

The *w*Mel gene WD0835 and its homologs encode a hypothetical protein that is conserved among *Wolbachia* strains in the first half of the protein while the C-terminal region is highly variable with numerous amino acid substitutions and indels, suggesting that the two regions evolve under different selective pressures (Figure 5A). In our experiment, WD0835 accumulated several non-synonymous mutations during passaging in Aa23 cells. In *w*Mel-infected cells, a SNP was detected in the variable region of the gene inducing a change in amino acid (Figure 5A). The SNP analysis indicated that the WD0835 mutant was at near fixation at 27 months post-transinfection, being present in ∼98% of the sequencing reads (Table S1). However, the SNP-specific qPCR data showed that the mutation, which started to increase in frequency about 20 months after transinfection, remained at frequencies below 50% (Figure 3C). This discrepancy between the sequencing and qPCR data is unlikely to be due to imperfect qPCR efficiency given that efficiencies where ∼100% both for the SNP-specific primers and the primers amplifying the control gene 16S to which it was compared (Figure S6A). The presence of within-sample polymorphism at the SNP position was also confirmed by Sanger sequencing with a double peak observed at 23, 24 and 27 months post-transinfection (Figure S5). One explanation could be that, for sequencing, a derived population of cells were maintained at high cell density in order to increase the *Wolbachia*-to-host DNA ratio, since this splitting regime was generally found to lead to higher symbiont densities (see methods, Figure S2). It is therefore possible that the mutation in WD0835 has a selective advantage that depends on *Wolbachia* density or the growth rate of host cells.

**Figure 5.**
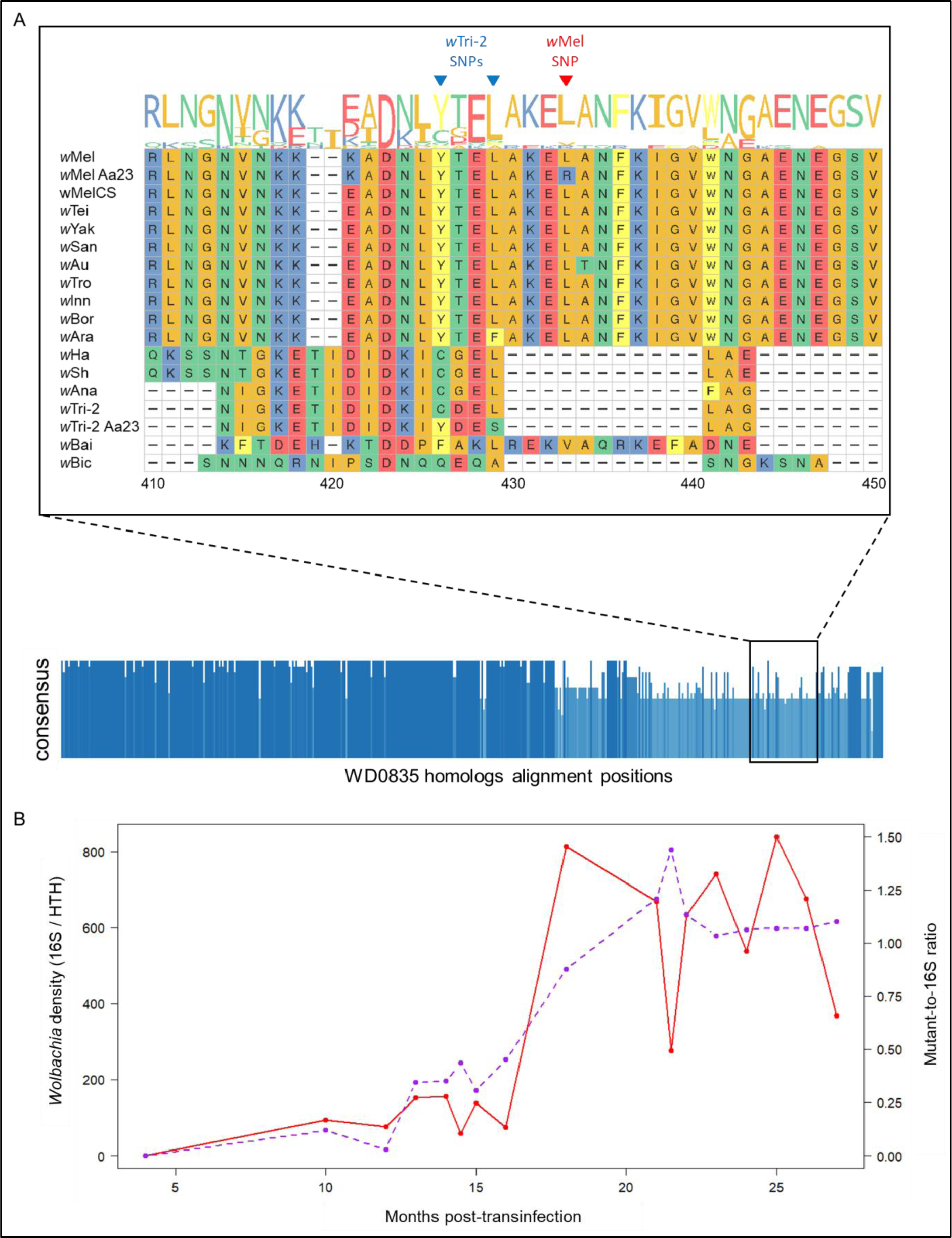
Evolution of WD0835 homologs. (A) Protein sequence alignment of WD0835 homologs visualized using the ggmsa visualization tool (http://yulab-smu.top/ggmsa/). Amino acids are coloured by chemistry in the zoom-in window (top); Blue bar chart: degree of sequence conservation across the protein alignment (bottom). (B) Variation in *w*Tri-2 density (red) and estimated frequency of the WD0835 homolog single/double mutant relative to 16S (purple).

Interestingly, in *w*Tri-2-infected cells, the homolog of WD0835 accumulated two non-synonymous SNPs in the same region of the gene as in *w*Mel (Figure 5A; Table S1). The first SNP was detected upon sequencing both 12 and 22 months post-transinfection, and the second SNP, located 9 nucleotides downstream, was only detected at month 22. Given the high proximity of the 2 SNPs on the gene, we could only design qPCR primers to distinguish between the double mutant and the ancestral variant, however, some cross amplification of the single mutant could not be excluded. Using these primers, we found an increase in the frequency of the single/double mutants from month 13 to fixation after 21 months. The spread of the mutant was strongly correlated with the sudden increase in *w*Tri-2 density in the cell line, suggesting a causal relationship between the two variables (Figure 5B).

### Evolution of a *PleD*-like response regulator

WD0221 is homolog of a *PleD* family two-component system response regulator which function in *Wolbachia* is unknown. In *w*Tro-infected Aa23 cells, a non-synonymous SNP changing an amino acid in the homolog of WD0221 appeared and spread between 10 and 15 months after transinfection and was maintained at high frequency to fixation (Figure 6A). Interestingly, the spread of this mutation also coincided with a global increase in symbiont density in the cell line suggesting that WD0221 may be involved in density regulation.

**Figure 6.**
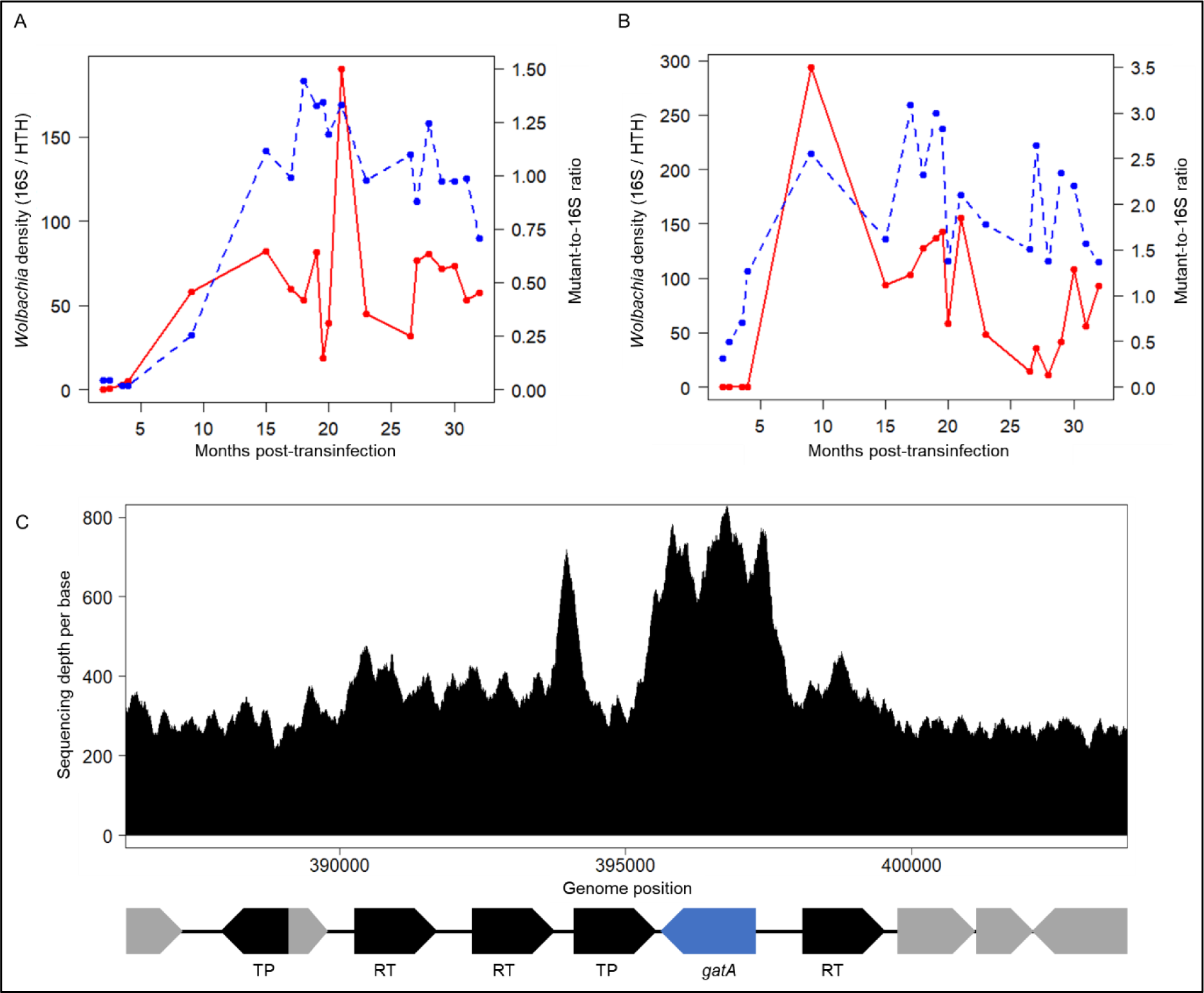
Evolution of WD0221 and WD0505 homologs. (A) Variation in *w*Tro density (red) and estimated frequency of the WD0221 homolog mutant relative to 16S (blue). (B) Variation in *w*Tei density (red) and estimated copy number of the WD0505 homolog relative to 16S (blue). (C). Sequencing depth of the region overlapping the *gatA*-encoding WD0505 homolog in the *w*Tei-infected Aa23 cell line. TP: transposase; RT: reverse transcriptase.

### Amplification of the amidotransferase *gatA*

WD0505 encodes a Glutamyl-tRNA(Gln) amidotransferase subunit A (*gatA*). In our experiment, we observed a ∼two-fold increase in sequencing depth in the region overlapping the homolog of WD0505 in *w*Tei-infected cells 17 months post-transinfection which we confirmed by qPCR, while the region was in one copy in *w*Tei-infected donor flies (Figure 6B and 6C, Figure S4 and S5B). In *w*Tei, *gatA* is flanked by transposable elements, which may have promoted the change in copy number, similar to what is observed for the Octomom region (Figure 6C). Monitoring of *gatA* copy number per *Wolbachia* genome following transinfection revealed that the amplification of the gene occurred and spread as early as 10 months post-transinfection, during which, a drastic increase in *w*Tei density was also observed (Figure 6B), supporting a potential role of the gene in density regulation. Copy numbers below 1 were observed at the beginning of passaging in Aa23 cells; however, we believe this could be explained by a drop in qPCR efficiency due to extremely low *w*Tei densities rather than a partial loss of the gene in the cell line.

## Discussion

Due to its obligate intracellular lifestyle, *Wolbachia* is not cultivable outside host cells which has greatly limited genetic studies. Cell culture has been widely used to investigate *Wolbachia*’s biology using transinfected or naturally-infected insect cell lines (O’Neill et al. 1997; Dobson et al. 2002; Furukawa et al. 2008; Xi et al. 2008; Raquin et al. 2015; Grobler et al. 2018; Fallon 2019; Fallon 2022). To our knowledge, only one study specifically looked at *Wolbachia* genomic changes during passaging in cell culture (Woolfit et al. 2013), however, this approach has not been leveraged as a tool to identify candidate genes associated with variation in symbiont density. Here, we generated new *Wolbachia*-infected mosquito cell lines and identified *de novo* mutations providing new insights into the evolution and genetic basis of symbiont density regulation.

### Gene function and putative role in density regulation

The Octomom region was consistently lost or maintained at very low frequencies in our cell lines, similar to what has been observed in previous cell culture studies (Woolfit et al. 2013; Grobler et al. 2018). Both the deletion and the amplification of Octomom are known to increase *Wolbachia* density in *Drosophila* hosts, however, the contribution of each of the eight Octomom-encoded genes in this effect is unclear (Chrostek et al. 2013; Chrostek and Teixeira 2015; Bénard et al. 2021; Duarte et al. 2021; Monnin et al. 2021). Some of the genes encode proteins with putative roles in DNA replication, transcription or repair and may directly affect bacterial replication (WD0507, WD0508, WD0509 and WD0510). Other Octomom genes harbour eukaryotic-like and/or toxin domains and could be bacterial effectors. WD0512, WD0513 and WD0514 are transcribed as part of an operon (Iturbe-Ormaetxe et al. 2005), have predicted outer-membrane localization, and distant homologues are present in the *Ae. aegypti* genome indicating horizontal gene transfer. WD0514 encodes ankyrin repeats involved in protein-protein interactions and believed to be important in *Wolbachia*-host interactions. WD0513 encodes Rhs repeats: in gram negative bacteria these have been associated with contact-dependent growth inhibition / mediation of intercellular competition (Koskiniemi et al. 2013), and can carry C-terminal toxin domains (Rhs-CT/WapA-CT) deployed to inhibit the growth of neighbouring cells. In *w*MelCS-infected cells, we also observed the spread of a mutation introducing a premature stop codon in the homolog of WD0508, suggesting that the gene has lost its function. This mutation was associated with a temporary but substantial rebound in Octomom copy number, indicating that it may have been adaptive. WD0508 harbours DNA-binding helix-turn-helix domains homologous to transcription regulators and might regulate the expression of other *Wolbachia* genes. Alternatively, WD0508 could be a secreted effector involved in *Wolbachia*-host interactions since its transgenic expression in *Drosophila* causes an increase in *Wolbachia* density (Lepage et al. 2017).

WD0835 homologs accumulated non-synonymous mutations in two of our mosquito cell lines which, in the case of *w*Tri-2, was associated with a strong shift in symbiont density. WD0835 encodes a hypothetical protein previously identified as a putative bacterial effector based on its phylogenetic distribution being restricted to the *Wolbachia* genus (Rice et al. 2017). Its conserved N-terminal region harbours structural homologies to various membrane and endoplasmic reticulum-associated proteins (HHpred probabilities >70% against PDBmm_CIF70_18_Jun database: Nucleoporin NSP1, Endoplasmic reticulum protein ERp29, Tail-anchored protein insertion receptor WRB) suggesting it may be involved in *Wolbachia*’s interactions with host membranes. Interestingly, WD0835 expression pattern throughout the host development is tightly correlated to that of a neighbouring gene, WD0830, with high levels of expression in *Drosophila* larvae and pupae (Gutzwiller et al. 2015; Rice et al. 2017). WD0830 encodes a protein named WalE1 shown to be a secreted effector that interacts with the host actin cytoskeleton and which heterologous expression increases symbiont densities (Sheehan et al. 2016; Martin and Newton 2023). Therefore, the WD0835 protein might act in concert with WalE1 to regulate symbiont proliferation through interactions with the host.

The homolog of WD0221 gained a non-synonymous mutation in the *w*Tro-infected cell line. WD0221 encodes the response regulator of a two-component PleC/PleD system conserved among *Wolbachia* strains (Christensen and Serbus 2015; Lindsey 2020). The PleC/PleD system is a widespread sensing and signalling pathway in bacteria that regulate many processes. While its role in *Wolbachia* has not been characterized, homologous systems in other intracellular bacteria are thought to play an important role in host cell infection (Kumagai et al. 2006; Lai et al. 2009).

WD0505 encodes the Glutamyl-tRNA(Gln) amidotransferase subunit A (GatA) of the GatCAB transamidosome complex, an enzyme that catalyses the transamidation of Glu-tRNA(Gln) in many bacteria, ensuring the fidelity of protein translation (Curnow et al. 1997). We observed the duplication of the *gatA* gene in *w*Tei-infected cells following transinfection which was associated with an increase in *w*Tei density. Thus, it is possible that this duplication caused an increase in the production of the GatA protein and as a consequence, had a general effect on protein synthesis and *Wolbachia* proliferation. As a support for the role of *gatA* in bacterial density, a *gatA* homolog in *Pseudomonas aeruginosa* was recently found to act as a secreted toxin that alters the stoichiometry of the GatCAB complex, leading to protein synthesis and growth defects in bacterial competitors (Nolan et al. 2021). Interestingly, (Woolfit et al. 2013) identified a mutation in WD0413, an aspartyl-tRNA synthetase (*aspC*), which is also involved in translation fidelity, during the passaging of *w*MelPop in mosquito cell lines. Thus, it is possible that the protein translation machinery is a primary target of selection during evolution in cell culture.

### Selection versus genetic drift in the cell culture environment

Similar to what was observed in (Woolfit et al. 2013), all the mutations we identified induced non-synonymous changes in *Wolbachia* genes suggesting they likely had phenotypic consequences. This is supported by the changes in symbiont density associated with most of these mutations. In addition, apart from the WD835 mutation in *w*Mel, all other mutations spread to fixation or near fixation, and some of them occurred independently in several cell lines, as was the case for the repeated loss of the Octomom region in *w*Mel, *w*MelCS and *w*MelPop. While we cannot exclude the role of genetic drift, these observations suggest that these evolutionary changes were more likely driven by selection. The complete absence of synonymous or intergenic mutations may be explained by a combination of low mutation rate (estimated to be ∼100-fold lower than the mitochondrial mutation rate at synonymous sites in *Drosophila* (Richardson et al. 2012)) and low recombination relative to the speed of a selective sweep when a beneficial mutation appears. Indeed, since *Wolbachia* reproduces clonally, the invasion of a mutant *Wolbachia* to fixation would inevitably erase any genomic diversity at other loci in the symbiont population.

While the identified mutations were most likely beneficial to the symbiont, in some cases we detected the presence of polymorphism for extended periods of time. For example, the Octomom region tended to be consistently lost but persisted at very low frequency in most replicates. This could be because selective pressures fluctuate in space or time in the cell culture environment. For instance, there is evidence that the Aa23 cell line is composed of different cell types with distinct morphologies (O’Neill et al. 1997). Moreover, competition for resources is likely to fluctuate rapidly in cell culture due to the alternation of host cell growth and stationary phases during passaging. Finally, temporal variation in host cell density may also determine the relative contribution of vertical and horizontal transmission of the symbiont. Therefore, the cell culture environment may allow the persistence of some polymorphism in the symbiont population if the fitness of different symbiont haplotypes varies with the host cell types, the level of competition within cells or the opportunities for horizontal transmission.

Evolutionary theory predicts that within hosts, overproliferating symbionts should have a competitive advantage and invade, even at the expense of the host’s fitness, a phenomenon called the “tragedy of the commons” (Hardin 1968; Frank 1996). In the case of vertically-transmitted symbionts, this may also increase the probability of being transmitted to the next generation. However, overproliferating mutants should be counter-selected at the between-host level if high symbiont density induces strong costs on the host fitness, as is the case in *Wolbachia* (Chrostek et al. 2013; Chrostek and Teixeira 2015; Martinez et al. 2015; Duarte et al. 2021). In Aa23 cell lines, high *Wolbachia* densities tended to be associated with slower cell growth and we would expect a similar trade-off on the evolution of symbiont density due to conflicting selective pressures within and between host cells. However, we also found that *Wolbachia* is transmitted horizontally between cells, which is in agreement with previous cell culture and insect studies (Frydman et al. 2006; White et al. 2017). If horizontal transmission is sufficiently high, host and symbiont fitness may be largely decoupled. Thus, costly overproliferating symbionts that slow the growth of host cells could still have an overall selective advantage in cell culture.

If the combination of vertical and horizontal transmission in cell culture is a potential explanation for the rapid evolution of symbiont density we observed, there are other non-mutually exclusive hypotheses. For instance, *Wolbachia* could have adapted in response to the change in host species or temperature following transinfection. Indeed, the symbiont strains used originated from a *Drosophila* host background maintained at 18°C for several years, while the Aa23 mosquito cell lines were kept at 28°C. However, in the case of the Octomom region, the loss of the region is probably independent of the host background or temperature since the same phenomenon was observed both in mosquito and *Drosophila* cell lines under different temperature regimes (this study and (Woolfit et al. 2013; Grobler et al. 2018)).

### Rate of evolution in cell culture versus the insect model

The mutations we characterized in our cell lines were all absent in the reference *Wolbachia* genomes sequenced from their natural *Drosophila* host species and from the *D. simulans* lines used for transinfection. This implies that these *Wolbachia* strains did not evolve over > 6 years spent in a new insect host background, whereas most of them accumulated mutations once maintained in cell culture over a much shorter timeframe. This is consistent with the relative stability of *Wolbachia* genomes observed following artificial host shifts or the introduction of the symbiont in the field for disease control applications (Huang et al. 2020; Morrow et al. 2020; Baião et al. 2021; Martinez et al. 2022). As discussed above, this may stem from different selective pressures between the cell culture and the insect models, in particular the fact that high-density symbionts may be rapidly eliminated from insect populations if they are too costly. A faster rate of evolution in cell culture may also be explained by larger population sizes and number of generations. Indeed, in insects, *Wolbachia* undergoes genetic bottlenecks at every host generation due to maternal transmission, giving less opportunities for new mutations to occur and be transmitted.

### Implications for disease control and future prospects

Whether *Wolbachia* genomes simply evolve faster in cell culture due to larger population sizes or because selective pressures are intrinsically different in this environment remains an open question. Further work is also needed to fully validate the link between the mutations we characterized and symbiont density, for example by conducting side-by-side comparison of the mutant phenotype to its ancestral variant. Nevertheless, our study demonstrates that the experimental evolution of *Wolbachia* strains in cell culture is a promising avenue for functional genetics in the absence of genetic transformation tools. Our results also have important implications for *Wolbachia*-based applications. Indeed, the long-term efficacy of disease control programmes depends on the stability of the symbiont’s capacity to invade mosquito populations and to inhibit virus replication (Vavre and Charlat 2012; Bull and Turelli 2013). Experimental evolution of *Wolbachia* strains could be used to generate new *Wolbachia* variants from cell culture showing interesting properties for disease control, followed by isolation and transfer into important disease vectors and evaluation for potential field releases. Finally, the cell culture model could be a powerful tool to study *Wolbachia*-host coevolutionary dynamics and underlying mechanisms. While we only investigated changes in the symbiont genome, we cannot rule out that some evolution in the host genome occurred in our cell lines, which could have had consequences on the observed changes in *Wolbachia* density.

## Methods

### *Wolbachia* strains, fly husbandry and transinfection protocol

The *Wolbachia* strains used for transinfection were sourced from the *D. simulans* STCP background (Table 1), except for *w*MelPop which was sourced from the *D. melanogaster* DrosDel *w*^1118^ isogenic background (Chrostek et al. 2013). All flies were maintained at 18°C on a cornmeal diet under a 12-hour light/dark cycle. Twenty-four hours before transinfection, a 96-well plate was seeded with *Wolbachia*-free Aa23 cells (∼1.6×10^5^ cells per well) in Schneider’s media (10% Fetal Bovine Serum, 1% PenStrep) and cells were left to attach to the bottom of the well overnight. On the day of transinfection and for each *Wolbachia* strain, *Wolbachia*-infected adult flies were allowed to lay eggs in BugDorm cages (W17.5 x D17.5 x H17.5 cm) with a Petri dish containing agar (3% agar, 1% sucrose, water) and a spot of yeast paste in the centre to stimulate egg-laying. After 1 hour, 500-1,000 fly embryos were collected from the Petri dish with a metal spatula, transferred to a 50 ml Falcon tube and rinsed in 50 ml of sterile water three times. Fly embryos were allowed to sink and the embryo pellet transferred to a 1.5 ml Eppendorf tube. Embryos were then surface-sterilized in 1 ml of 50% commercial bleach solution for 2 min and rinsed in 1 ml of 70% ethanol for 5 min twice and finally 1 ml of sterile water. Embryos were then resuspended in 1 ml of Schneider’s media and homogenized with a sterile pestle. The embryo homogenate was centrifuged at 2,500 g for 10 min to remove eggshells and cell debris. 100 µl of the supernatant was overlaid onto Aa23 cells. When fully confluent, mosquito cells were serially passaged from the 96-well plate, to 48-, 24- and 12-well plates. Cells were later maintained in 25 cm^2^ flasks in Schneider’s media at 28℃ and transferred to new flasks every 7-10 days by diluting 1 mL of cell suspension into 4 mL of fresh media (1 in 5 dilution).

### Quantification of *Wolbachia* density

For each sample, DNA was extracted from ∼1.6×10^6^ Aa23 cells resuspended in 200 μL of STE buffer followed by a 30 min incubation at 65°C with 2 μL of Proteinase K (20 mg/mL) and a final incubation for 10 min at 95°C. For DNA extraction from flies, the same was done by homogenizing a single adult female in STE buffer with a sterile pestle. The extracted DNA was then diluted 1/5 in water and samples were centrifuged at 2,000 g for 2 min before endpoint PCR or qPCR.

*Wolbachia* density relative to host DNA was measured by qPCR using primers amplifying the single copy *Wolbachia* 16S rRNA gene and the mosquito Homothorax gene (HTH) as a control (Table S2). For each gene, 2 μL of DNA template and the Fast SYBR™ Green Master Mix (Applied Biosystems) were used in 10 μL reactions: 5 μL of master mix, 2 μL of water, 0.5 μL of each 5 μM primer and the following cycles: 95°C for 20 s, 40 cycles of 95°C for 1 s and 60°C for 20 s, followed by a melt-curve analysis. For each sample and each gene, two qPCR reactions were done. *Wolbachia* density was then estimated by averaging the Cq values of the two technical replicates per gene and the following formula: *2^(meanHTH Cq - mean 16S Cq^*^)^.

### In situ detection of Wolbachia

Cells were plated out overnight in a poly-L-lysine coated 24-well glass-bottom plate (Corning, New York, USA) at a density of 5×10^5^ cells per well. Schneider’s media was then removed and cells were fixed with 200 µL of ice cold 10% formalin for 30 minutes at room temperature. Cells were washed twice in PBST (PBS, 0.1% Tween^TM^ 20) then incubated overnight at 37°C in a hybridisation buffer containing 50% formamide, 25% 20X SSC, 0.2% dextran sulphate, 2.5% herring sperm DNA, 1% tRNA, 1% Denhardt’s solution, 0.015% DTT and 100 ng/ml of each *Wolbachia* 16S rRNA probe. To detect all transinfected symbiont strains, the following *Wolbachia* probes were combined: W2 5’- CTTCTGTGAGTACCGTCATTATC-(Cyanine3)-3’, W3 5’- AACCGACCCTATCCCTTCGAATA-(Cyanine3)-3’ (Moreira et al. 2009). In order to distinguish the transinfected Supergroup A strains (*w*Au and *w*Mel) from *w*AlbB in the co-culture experiment, we combined a Supergroup A-specific probe (5’- ACCTGTGTGAAACCCGGACGAAC-(Alexa fluor 488)-3’) with a *w*AlbB-specific probe (5’- TAGGCTTGCGCACCTTGCAGC-(Cyanine3)-3’). Following this, samples were washed twice in a solution with 5% 20X SSC, 0.015% DTT and twice in 2.5% 20X SSC, 0.015% DTT for 20 minutes per wash at 55°C. Cells were then stained for 20 minutes with Hoechst 33342 (ThermoFisher, MA, USA) at a concentration of 1 µg/mL in PBS. Finally, cells were washed with PBS and left in PBS for visualisation to prevent drying out. All samples were visualised on a Zeiss LSM 880 confocal microscope (Zeiss, Oberkochen, Germany).

### *Wolbachia* purification and DNA preparation for sequencing

To ensure enough DNA material for sequencing, a population of Aa23 cells was derived from each of the *Wolbachia*-infected cell lines and maintained in Schneider’s media in two 225 cm2 flasks per symbiont strain. In order to increase the *Wolbachia*-to-host DNA ratio, cells were passaged twice in 225 cm2 flasks every 10 days using a 1 in 2 dilution, instead of 1 in 5, as higher cell density was found to increase *Wolbachia* density (Figure S2). Cells from the two flasks were then harvested and concentrated by centrifugation at 2,500 g for 2 min. The cell pellets were pooled, resuspended in 2 mL of Schneider’s media and homogenized with 1mm beads in 2 mL screw cap tubes in a tissue lyzer. The homogenate was then centrifuged at 2,000 g for 2 min to remove cellular debris and the supernatant passed through 5 and 2.7 µm pore size filters. The filtrate was then centrifuged at 18,500 g for 15 min to pellet the bacteria. The supernatant was discarded and the bacterial pellet resuspended in 1 mL of Schneider’s media and the previous centrifugation step repeated once. The final resuspended bacterial pellet was then incubated with 20 µL of DNase I at 37°C for 30 min to digest remaining host DNA. Following digestion, the sample was centrifuged at 18,500 g and the supernatant discarded. DNase I was inactivated by incubating the sample at 75°C degrees for 10 min. Finally, DNA was extracted with the Gentra Puregene tissue kit (Qiagen) and the DNA pellet resuspended in 100 µL of nuclease-free water.

### Illumina sequencing, variant and sequencing depth analysis

DNA libraries were prepared using the Kapa LTP Library Preparation Kit (KAPA Biosystems, Roche7961880001) and sequenced on the Illumina MiSeq platform with the MiSeq Reagent Kit v3 to generate 2×150 bp reads. Raw reads were demultiplexed using bcl2fastq and adapters were trimmed with Trimmomatic v0.38.0 (Bolger et al. 2014). Host reads were filtered out by mapping the Illumina reads against the *Ae. albopictus* reference assembly (Genbank accession: GCA_006496715.1) using BWA-MEM (Li 2013). Unmapped reads were then mapped to their respective *Wolbachia* reference genome (Table 1) with BWA-MEM. Reads mapped onto the symbiont genome were filtered with bamtools v2.5.2 (Barnett et al. 2011) to remove duplicates and retain properly paired reads with a minimum mapping quality of 20 (MAPQ), except for reads with a MAPQ score of 0 in order to keep reads mapping to repetitive regions. The SNP analysis was then conducted using the Snippy pipeline v4.6.0 (Seeman, Torsten. Snippy: fast bacterial variant calling from NGS reads; 2020 https://github.com/tseemann/snippy). SNPs were called with a threshold of 10 reads minimum coverage, 0.9 minimum proportion, >30 Phred quality scores for variant evidence (i.e. 99.9% accuracy). Detected variants were visually inspected by loading the BAM alignment files into the Artemis genome viewer (Carver et al. 2012). Finally, sequencing depth per base was calculated for each BAM file with the samtools depth command (Li et al. 2009) and visualised with the R software (R Core Team 2013).

### Timecourse analysis of mutations

SNPs and short indels were first validated by Sanger sequencing using PCR products generated with primers listed in Table S2. The timecourse analysis of the identified mutations was then conducted using qPCR and SNP-specific primers. To this end, primer pairs were designed with one of the primers having the last or the penultimate base corresponding to the mutated allele on the SNP (Table S2). PCR efficiencies were assessed with a dilution series (Figure S6A) and the specificity of the amplification was validated by ensuring that no or negligible amplification was observed for samples carrying the ancestral allele (Figure S6B). The copy number of the target SNP relative to 16S was calculated from the average of two technical replicates per sample as follows: *2^(mean 16S Cq – mean SNP Cq)^*. Relative SNP copy number was then used as a proxy for the frequency of the SNP in the symbiont population. Copy number variation for the Octomom region (WD0513) and *gatA* (WD0505) were calculated in the same way.

In order to test for within-sample polymorphism in Octomom copy number, endpoint PCR with primers amplifying the WD0513 gene and primers flanking the Octomom region was conducted to detect the Octomom-carrying and deleted variants respectively (Table S2). The *wsp* gene was used as a control for *Wolbachia* presence (Table S2). Each set of primers was used in separate reaction with 2 μL of DNA template and the 2X DreamTaq Green PCR Master Mix (ThermoFisher) in a 21 μL reaction: 9.4 μL of master mix, 9.4 μL of water, 0.1 μL of each 20 μM primer. The following PCR cycle was used: 95°C for 3 min, 35 cycles of 30s denaturation at 95°C, 30s for primer annealing at 56 °C, 3 min extension at 72°C and a 15 min final extension step at 72°C.

## Data Availability

Raw sequencing reads are available at the Sequence Read Archive under the following accession: TBA.

## Acknowledgements

This work was supported by the Wellcome Trust (grant numbers 202888/Z/16/Z and 226166/Z/22/Z).

---

**Figure S1.**
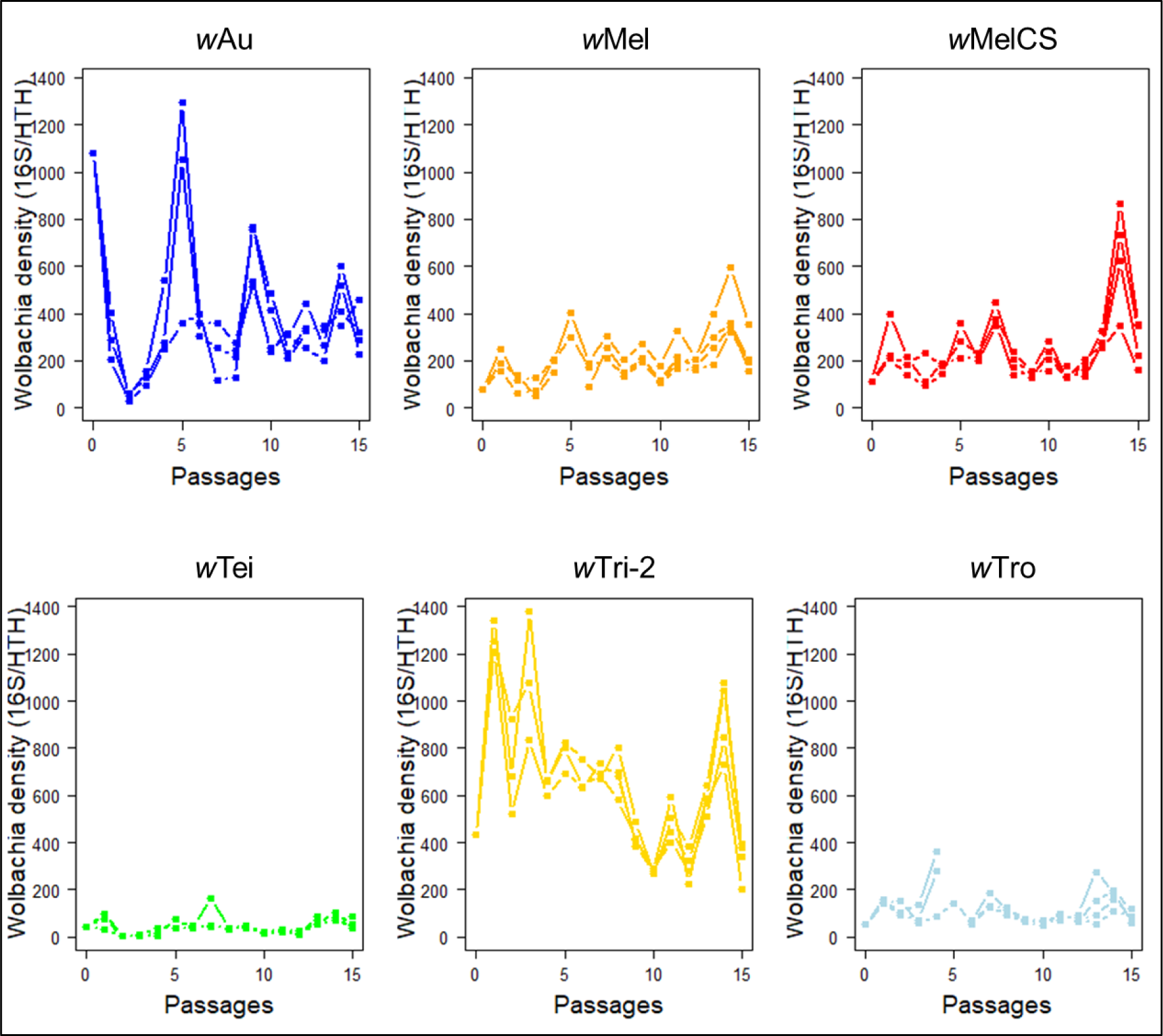
Short-term dynamics in *Wolbachia* density. For each *Wolbachia* strain, 3 replicates cell lines were derived from the same original flask 27 months post-transinfection (22 months for *w*Tri-2) and maintained in parallel. Density was measured at each passage into new flasks every 10 days.

**Figure S2.**
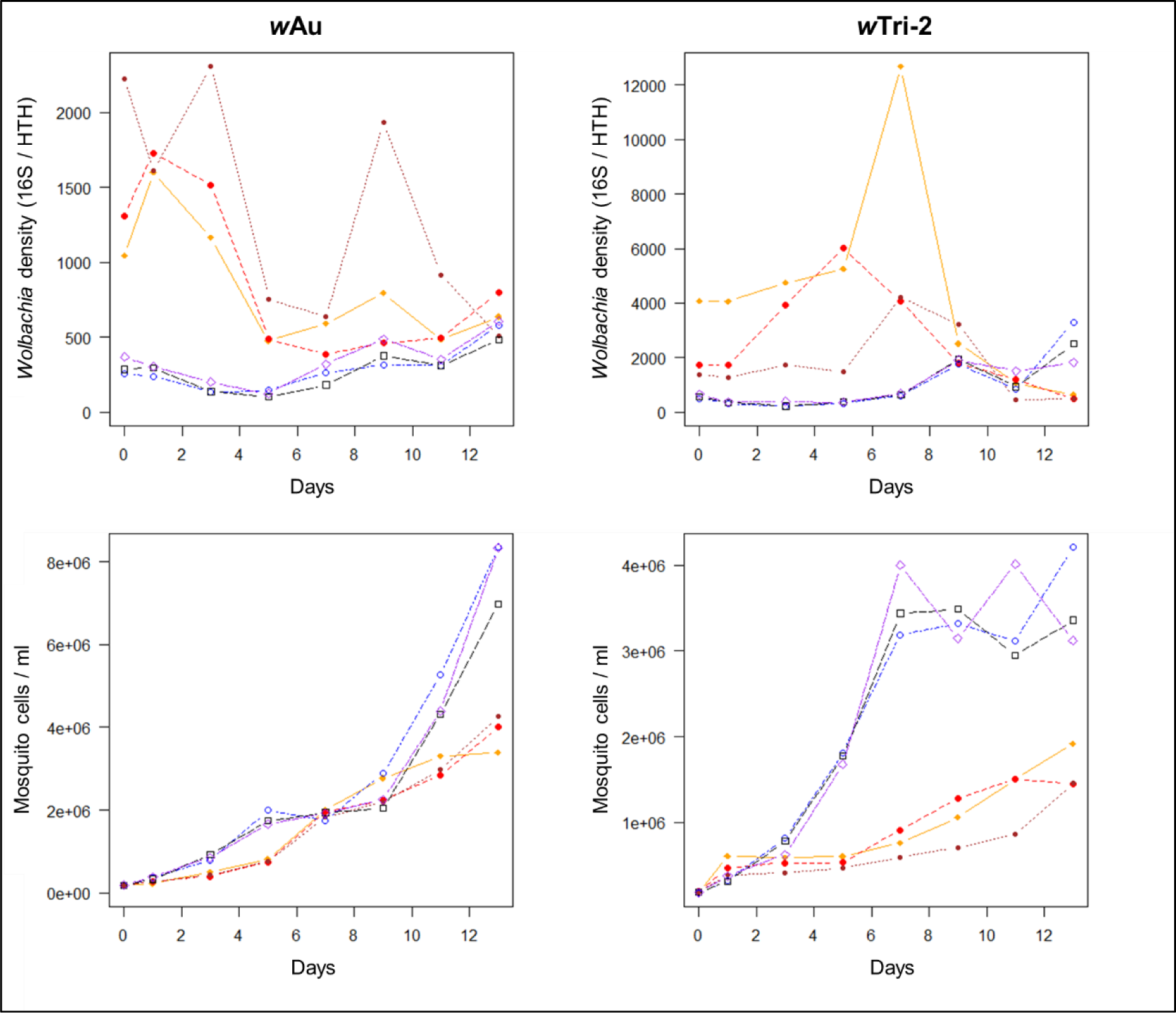
Relationship between *Wolbachia* density and host cell growth rate. Six cell lines originating from the same flasks were derived from either *w*Au- (left) or *w*Tri-2-infected (right) mosquito cells. Three replicate lines per *Wolbachia* strain were first passaged at low cell density (∼8×10^5^ cells: blue, black, purple) and the three other lines at high cell density (∼1.6×10^6^ cells: orange, red, brown) for four passages before the start of the experiment. Several flasks per replicate cell line were then seeded with 1.6×10^6^ cells and *Wolbachia* density (top) and mosquito cell concentration (bottom) were measured every two days (each datapoint corresponds to a different flask).

**Figure S3.**
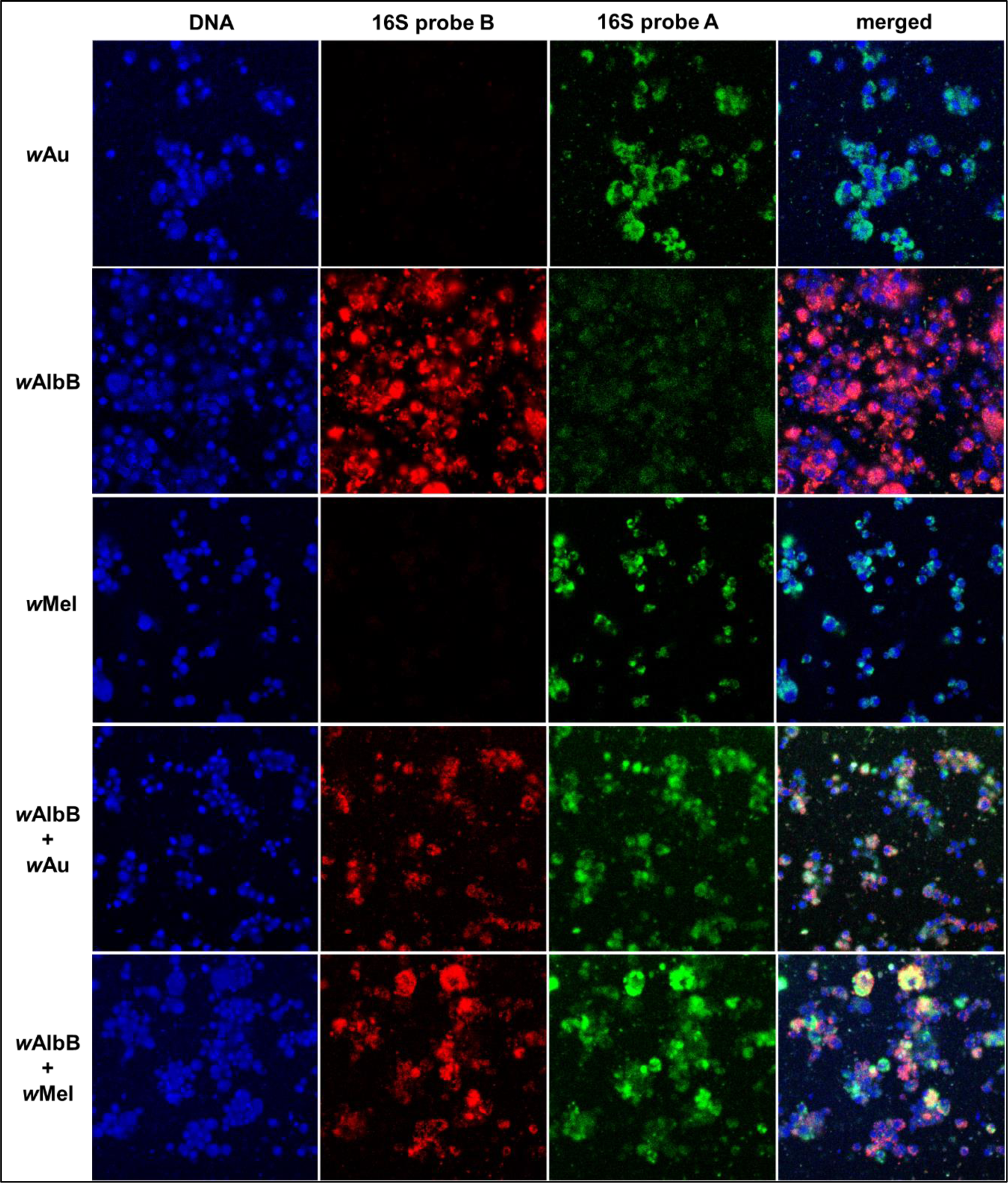
Fluorescent *in situ* hybridization imaging of *Wolbachia* strains in single or co-culture in Aa23 cells. Cells were stained with Hoechst 33342 (DNA, blue), 16S rRNA probe A (*w*Au/*w*Mel-specific, green) and probe B (*w*AlbB-specific, red).

**Figure S4.**
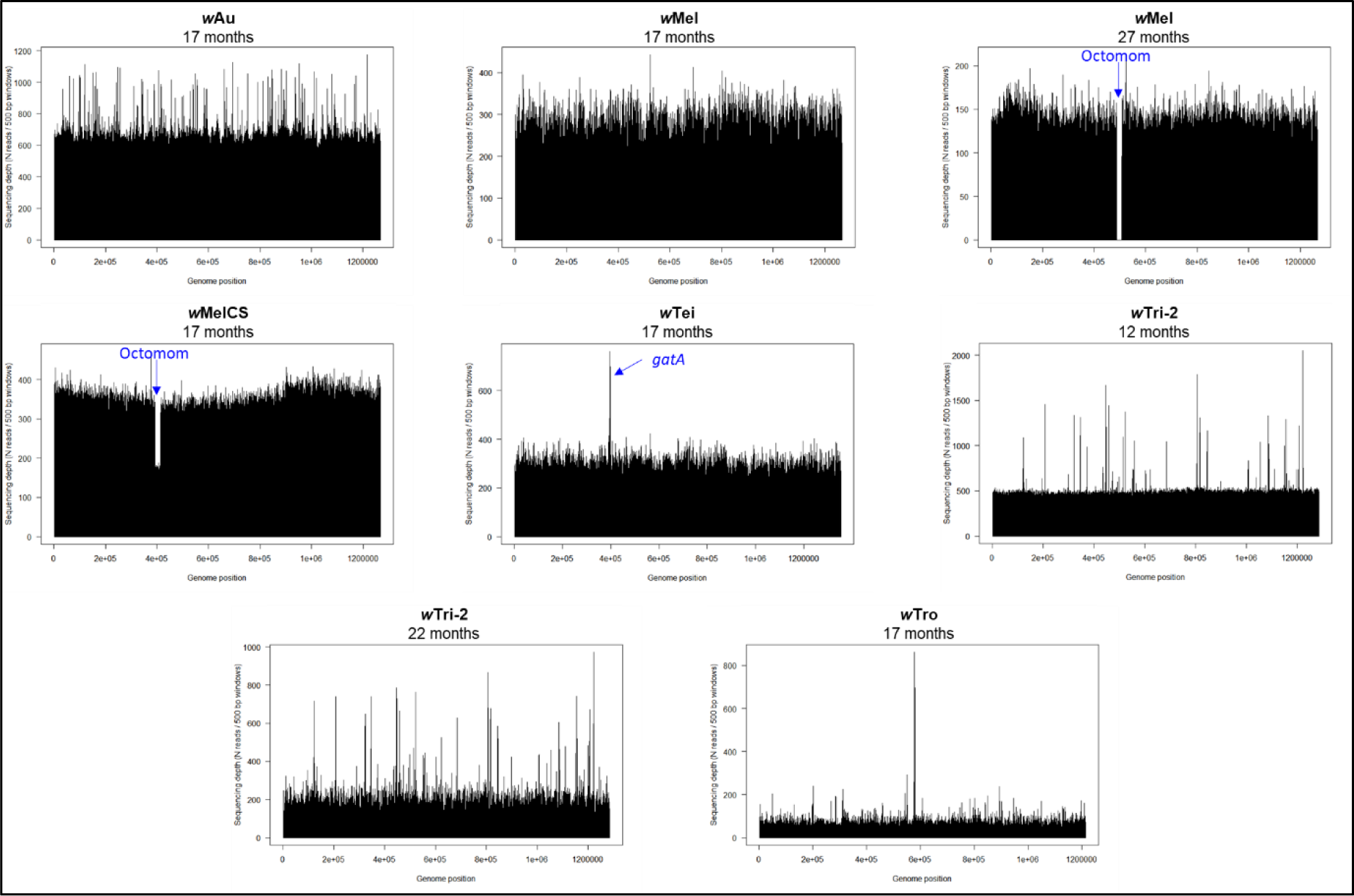
Sequencing depth of *Wolbachia* genomes in Aa23 cells. Sequencing depth was averaged over 500 bp windows.

**Figure S5.**
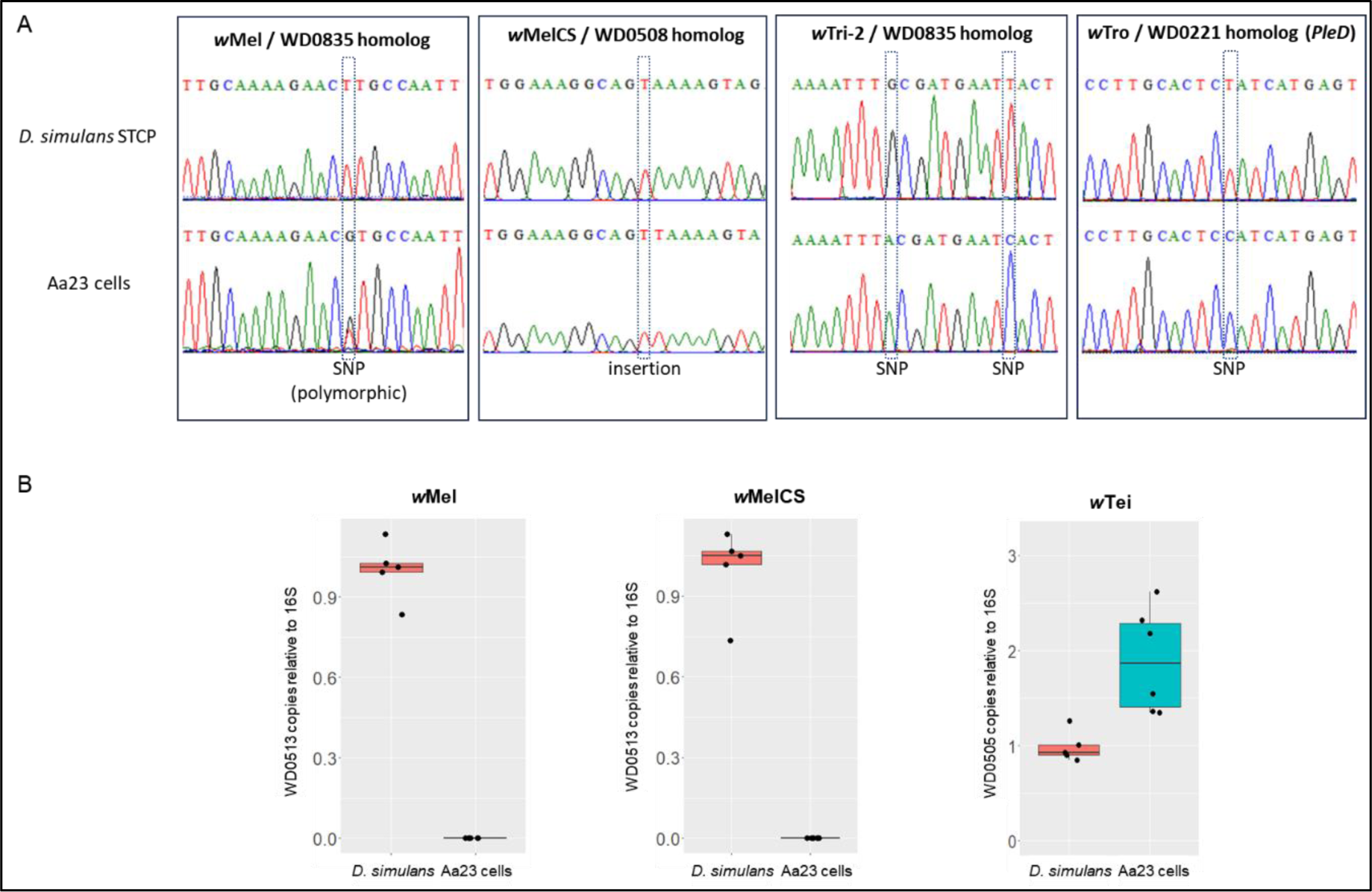
Validation of detected mutations in *Wolbachia* genomes. (A) Sanger sequencing chromatograms of SNPs and small indels identified in transinfected *Wolbachia* strains in *D. simulans* and their Aa23 cell line counterpart. (B) qPCR validation of copy number variation in Octomom region (WD0513) and the *w*Tei homolog of WD0505. Single adult females were used for *Wolbachia*-infected *D. simulans* donor lines. Samples collected after 27 months post-transinfection are shown for *Wolbachia*-infected Aa23 cells.

**Figure S6.**
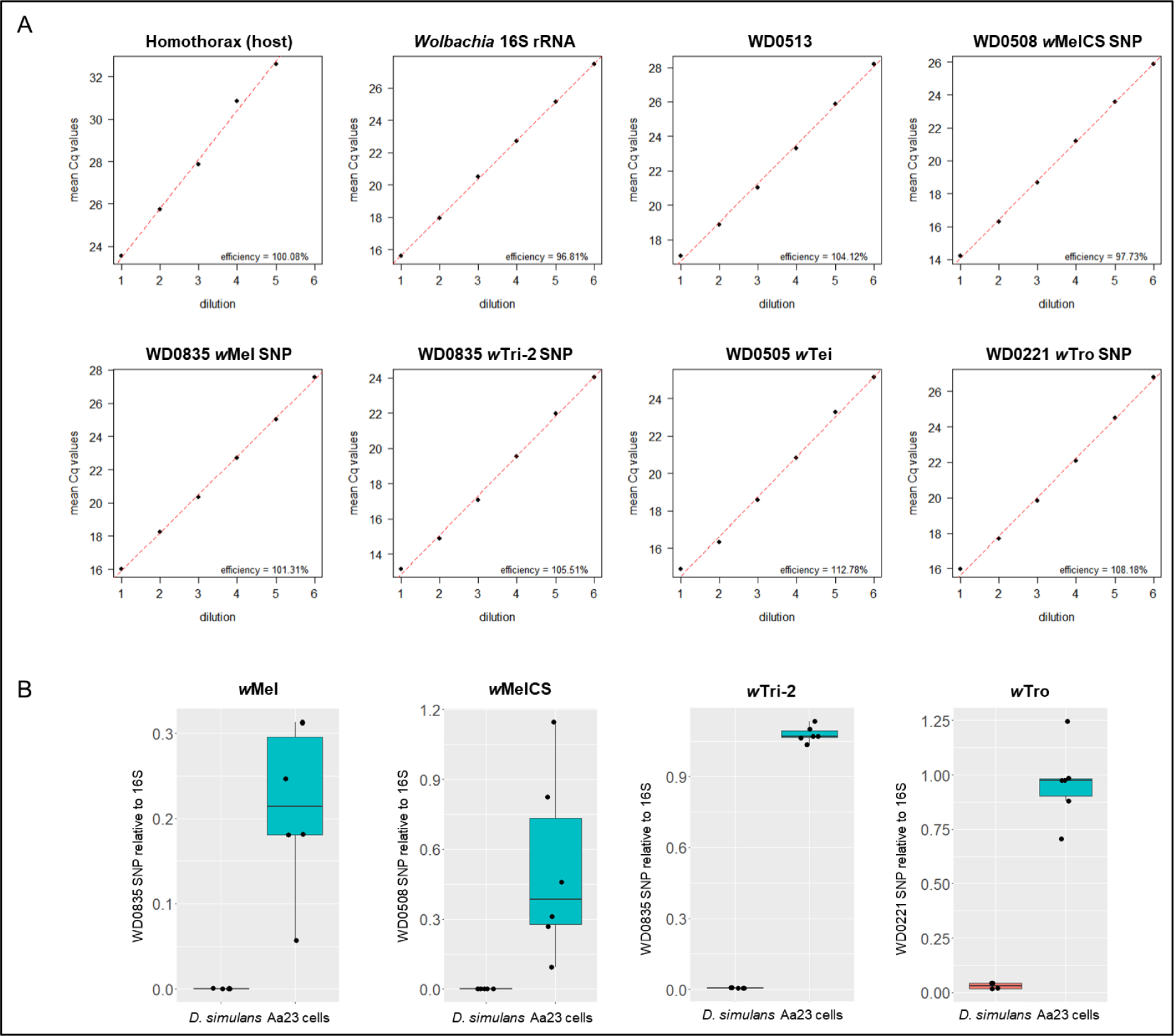
Efficiencies and specificity of qPCR primers used. (A) 1 in 5 dilution series. Each datapoint is the mean of two technical replicates. Red line: regression model used to estimate qPCR efficiencies of each primer pair. (B) qPCR validation of SNP-specific primer specificity. Single adult females were used for *Wolbachia*-infected *D. simulans* donor lines. Samples collected after 27 months (22 months for *w*Tri-2) post-transinfection are shown for *Wolbachia*-infected Aa23 cells, except for the WD0508 SNP in *w*MelCS for which samples collected between 15-20 months post-transinfection are shown.

